# A prospectively validated machine learning model for the prediction of survival and tumor subtype in pancreatic ductal adenocarcinoma

**DOI:** 10.1101/643809

**Authors:** Georgios Kaissis, Sebastian Ziegelmayer, Fabian Lohöfer, Hana Algül, Matthias Eiber, Wilko Weichert, Roland Schmid, Helmut Friess, Ernst Rummeny, Donna Ankerst, Jens Siveke, Rickmer Braren

## Abstract

**Purpose:** To develop a supervised machine learning algorithm capable of predicting above vs. below-median overall survival from medical imaging-derived radiomic features in a cohort of patients with pancreatic ductal adenocarcinoma (PDAC).

**Materials and Methods:** 102 patients with histopathologically proven PDAC were retrospectively assessed as the training cohort and 30 prospectively enrolled patients served as the external validation cohort. Tumors were segmented in pre-operative diffusion weighted-(DW)-MRI derived ADC maps and radiomic features were extracted. A *Random Forest* machine learning algorithm was fit to the training cohort and tested in the external validation cohort. The histopathological subtype of the tumor samples was assessed by immunohistochemistry in 21/30 patients of the external validation cohort. Individual radiomic feature importance was evaluated.

**Results:** The machine learning algorithm achieved a sensitivity of 87% and a specificity of 80% (ROC-AUC 90%) for the prediction of above- vs. below-median survival on the unseen data of the external validation cohort. Heterogeneity-related features were highly ranked by the model. Of the 21 patients for whom the histopathological subtype was determined, 8/9 patients predicted by the model to experience below-median overall survival exhibited the quasi-mesenchymal subtype, while 11/12 patients predicted to experience above-median survival exhibited a non-quasi-mesenchymal subtype (Fisher’s exact test P<0.001).

**Conclusion:** The application of machine-learning to the radiomic analysis of DW-MRI-derived ADC maps allowed the prediction of overall survival with high diagnostic accuracy in a prospectively collected cohort. The high overlap of clinically relevant histopathological subtypes with model predictions underlines the potential of quantitative imaging workflows in pre-operative subtyping and risk assessment in PDAC.

## Introduction

Pancreatic ductal adenocarcinoma (PDAC) carries among the poorest prognoses of all cancers. Tumors exhibit significant heterogeneity on a genetic, transcriptomic and proteomic level, which manifests itself in a complex tissue architecture including tumor cells, various fibroblast and immune cell populations embedded in a poorly vascularized, dense stroma (1). Despite its overall dismal prognosis, recent research has identified specific molecular subtypes with distinct therapy response and outcome. Among these, the so-called classical phenotype shows improved chemotherapy response and survival compared to the so-called quasi-mesenchymal or basal-like subtype underlining the urgent requirement for advanced techniques for precise pretherapeutic patient stratification (2,3). This is key to both, adequate patient management, based on informed decision processes, clinical trial design and outcome interpretation.

In heterogeneous tumors such as PDAC, biopsies carry a significant risk of tissue undersampling. In contrast, imaging inhabits a unique niche in precision medicine in that it can provide volumetric information non-invasively. Radiomics, the process of derivation of quantitative analytics from medical imaging data (4) represents a significant advance over traditional image-analysis workflows as it leverages data science and machine learning techniques to exploit non-intuitive image content and integrate it with clinical information to create a generalizable model capable of predicting e.g. biological features or the course of disease (5).

Since PDAC is a relatively rare tumor entity, typically only treated in specialized interdisciplinary centers, there is still a paucity of radiomic studies aiming to assess pertinent metrics such as patient survival or histopathological subtypes. Our study aims to contribute to this field by applying a standardized, reproducible radiomic workflow pipelined to an efficient and explainable machine learning model capable of predicting overall survival and showing highly significant correlation with relevant histopathological subtypes, retrospectively trained and prospectively validated on a cohort of PDAC patients.

## Methods

### Study design

Data collection, processing and analysis were approved by the institutional ethics committee (Ethics Commission of the Faculty of Medicine, protocol numbers 180/17 and 5573/12). The study was designed as a retrospective cohort study with a prospective validation cohort. The requirement for written consent was waived for the retrospective cohort and written consent was obtained for the prospective cohort. All procedures were carried out in accordance to pertinent laws and regulations.

We considered patients with final histopathological diagnosis of PDAC of the head and body for inclusion in the study. Patients who did not have a final diagnosis of PDAC, had undergone treatment such as chemotherapy or resection prior to enrolment, refused treatment or study inclusion, died within the first 2 months of follow-up (to limit bias from postoperative complications), did not undergo the full imaging protocol or did not have technically sufficient imaging available, were excluded. For inclusion in the training cohort, we retrospectively considered 206 consecutive patients, who presented at our institution between 2008 and 2013 and underwent imaging at the department of radiology with a suspected finding of PDAC. The follow-up interval was defined as 5 years post-imaging. Follow-up was handled by the departments of surgery and internal medicine. A total of 102 patients were included in the study as the training cohort.

Prospective patients were accrued from 2013 onwards. Participants underwent PET/MRI evaluation at the time of diagnosis. Of 62 consecutive patients who were considered for inclusion, 30 patients fulfilled the enrolment criteria and designated as the external validation cohort.

Clinical data was sourced from the clinical information system. Radiomics data was generated during data analysis. For exclusion of bias, data analysis was performed in pseudonymized form and handled by separate individuals (G.K. and S.Z.). Data analysis was performed starting in June 2018. Patient flowcharts and the complete STROBE checklist can be found in the supplementary material.

### Clinical variables

The following clinical data was collected for patients in the training and external validation cohorts: age at diagnosis, sex, p/cTNM, R, G, tumor volume (from the final histopathological report or calculated from the segmentation volume), ECOG-state and chemotherapy regimen. Where applicable and available, pre-operative CA19-9 levels and lymph-node ratios (LNR) were noted. Overall survival was defined as the time from diagnosis to disease-related death.

### Imaging data acquisition

The 102 training cohort patients underwent magnetic resonance imaging (MRI) at 1.5T (Siemens Magnetom Avanto, release VB17). The protocol included the following sequences: axial and coronal T2-weighted spin echo (SE) images at 5mm; axial T1w gradient echo (GE) images at 5mm before contrast media injection and during the arterial, pancreatic parenchymal, portal-venous, systemic venous and delayed phases (as determined by testing bolus injection); axial unidirectional diffusion-weighed imaging at b-values of 0, 50, 300 and 600 with echo-planar imaging (EPI) readout and ADC map calculation. ADC map reconstructions were 5.5×5.5×5 mm (xyz) to a 192×192 voxel matrix. Furthermore, single-shot T2w magnetic resonance cholangiopancreatography (MRCP) was performed and reconstructed as a radial maximum intensity projection (MIP) series. The 30 external validation cohort patients underwent MRI on a 3T clinical PET-MRI scanner (Siemens Biograph mMR, VB18) at the hospital’s nuclear medicine department. The protocol was performed as above with the following alterations: ADC-map reconstructions were 5.1×5.1×5.1 mm (xyz) to a matrix of 192×192 voxels; furthermore an axial spectral adiabatic inversion recovery (SPAIR) fat-suppressed post-contrast sequence at 5 mm and a whole-body positron emission tomography scan after application of 18F-Fluordesoxyglucose (FDG) were included. The imaging protocols used, and the technical hardware specifications of the MRI machines remained unaltered during the data acquisition period.

### Data segmentation

Pseudonymized datasets were exported from the hospital picture archiving system (PACS) onto a radiological workstation and segmented under reporting room conditions by consensus reading of 2 experienced observers (G.K. and S.Z.) and quality-controlled by an abdominal radiologist with >10 years of experience in pancreatic MRI (R.B.). Segmentation was performed manually in the b=600 images and transferred to the ADC maps. All other sequences were available to observers for anatomical correlation.

### Inferential statistical modeling

For assessing potential clinical confounding parameters introducing bias to the survival prediction, survival time was modeled in both cohorts using a multivariate *Cox proportional hazards* model. The distributions of clinical variables were compared between groups using *Fisher’s exact test*. For subsequent machine learning modeling, the two cohorts were dichotomized by median overall survival to yield two sub-cohorts of equal size. ROC-thresholds were evaluated with the *Kolmogorov-Smirnov-*statistic. Biostatistical modeling was performed in SPSS Version 25. For all inferential statistical procedures, a P-value of <0.05 was considered statistically significant.

### Image postprocessing, radiomic feature extraction and machine learning modeling

All steps of image postprocessing, feature extraction, feature preprocessing, feature engineering and machine learning modeling are detailed in the supplemental material. In brief, radiomic features were derived using *PyRadiomics* (v. 2.1 (6)) yielding a total of 1688 features, of which 504 were retained after preprocessing. A *Random Forest* (7) classifier was fit in a supervised fashion with survival above versus below median serving as label to the training cohort radiomic features and tested for predictive sensitivity, specificity and ROC-AUC in the external validation cohort. Feature importance was assessed to derive significant radiomic features for the model. All analyses were carried out using the *Python* programming language.

### Histopathological workup of tumor samples

Histopathological staining and immunohistochemical workup were performed as described in (8). In brief, staining for the markers *HNF1a* and *KRT81* was carried out and tumors categorized into three subtypes: *classical, exocrine* and *quasi-mesenchymal*. Tumors not positive for either marker were designated *unclassifiable. Classical, exocrine* and *unclassifiable* tumors are onwards referred to as *non-quasi-mesenchymal*.

## Results

The distribution of clinical parameters did not differ significantly between the training and the external validation cohorts. Among the clinical parameters, the choice of chemotherapy regimen (gemcitabine vs. FOLFIRINOX) was significantly associated with overall survival in the training cohort but not in the testing cohort and the percentage of patients receiving each regimen was identical (with ∼70% of patients receiving gemcitabine in each cohort, *Fisher’s exact test* P=1.0). Metastatic status at baseline was significantly associated with diminished survival in both cohorts and was also identically distributed (∼25% of patients, *Fisher’s exact test* P=.81) in both cohorts. The results of multivariate Cox analysis and crosstabulations can be found in the supplementary material.

The Random Forest algorithm achieved a sensitivity of 86.7%, a specificity of 80.0%, a positive predictive value of 81,2% and a negative predictive value of 85,7% (*Fisher’s exact test* P=0.0007). The area under the ROC curve calculated on the external validation cohort data was 0.90 (Fig. 1). Further model evaluation metrics can be found in the supplementary material.

**Fig. 1:**
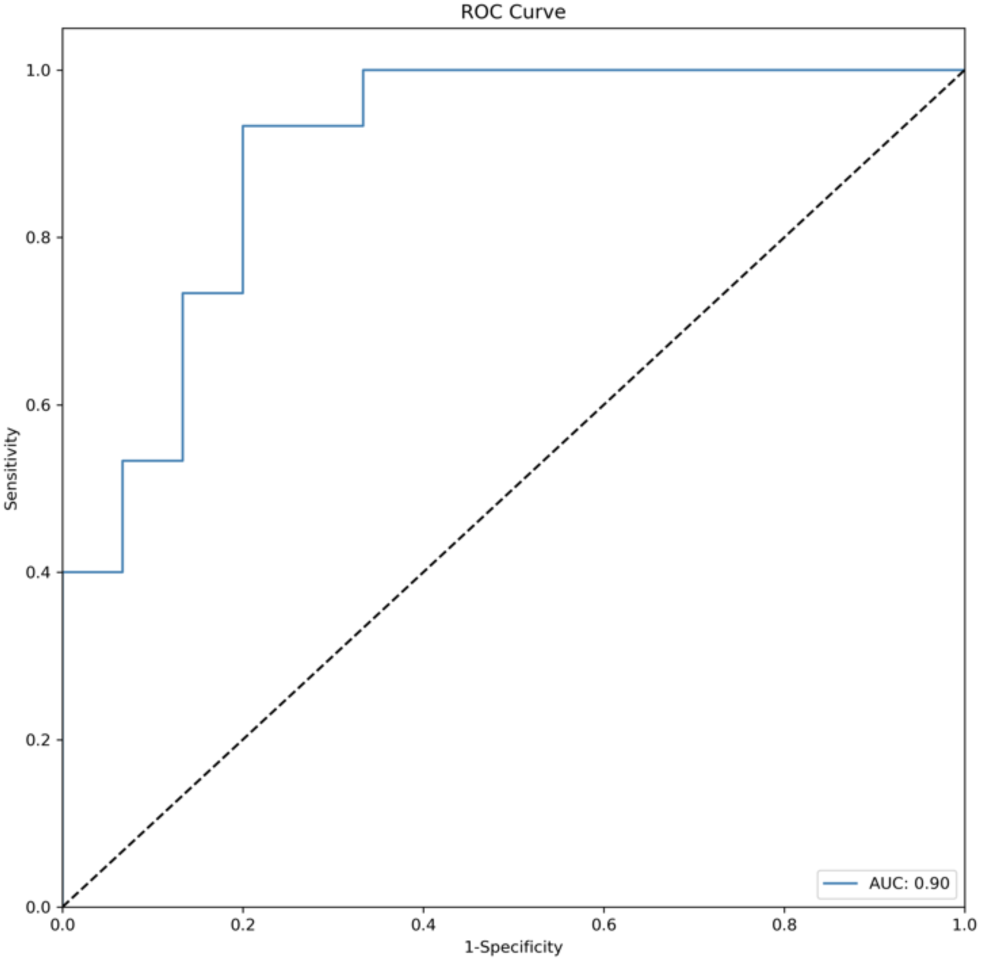
ROC curve of model performance on the external validation cohort. The classification threshold was 0.5, resulting in a ROC-AUC of 0.9. N=30 patients.

Furthermore, the algorithm’s predictions enabled statistically significant stratification of above- vs. below-median overall survival in the external validation cohort (*log-rank-test* P <0.001, predicted median survival for the below-median 17.0 months vs. 31.3 months for the above-median group) with the resulting predicted survival curves showing near-perfect overlap with the actual survival times of the patients (Fig. 2).

**Fig. 2:**
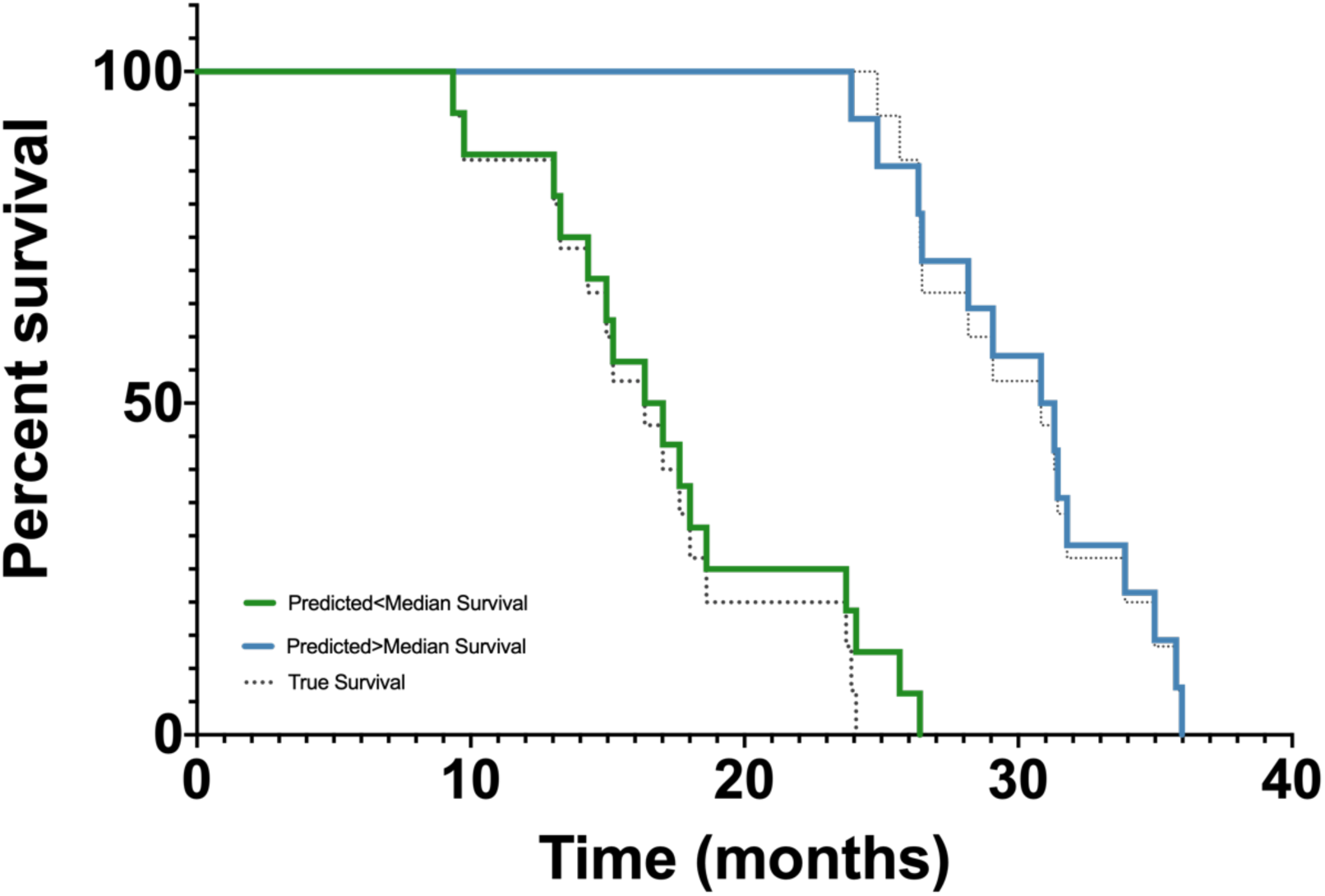
Kaplan Meier curves showing the predicted survival (blue and green curves) and the true survival (dotted curves) for patients in the external validation cohort. Log-rank-test between predicted survival curves: P<0.001, N=30 patients total.

The histopathologic subtype of the tumor samples could be determined for 21 of the 30 patients in the external validation cohort. The *quasi-mesenchymal* histopathological tumor subtype was greatly overrepresented in the patient collective predicted by the algorithm to experience below-median survival with 8 out of 9 patients having *quasi-mesenchymal* subtype tumors. The opposite also held true, with 11 out of 12 patients predicted by the algorithm to experience above-median survival having *non-quasi-mesenchymal* subtype tumors (Fisher’s exact test P<0.001, Table 1).

**Table 1:** Overlap between predicted survival groups and histopathological subtypes. The *quasi-mesenchymal* subtype was highly overrepresented in the group with predicted below-median survival and the *non-quasi-mesenchymal* subtypes in the group with predicted above-median survival. Fisher’s exact test P<0.001, N=30 patients.

|  | *Quasi-mesenchymal*<br>*subtype* | *Non-quasi-mesenchymal*<br>*subtype* |
| --- | --- | --- |
| *Predicted survival >Median* | 1/12 (9%) | **11/12 (91%)** |
| *Predicted survival <Median* | **8/9 (89%)** | 1/9 (11%) |

Feature importance evaluation yielded 8 highly important features (Fig. 3). Seven of these features (*GLCM Difference Variance, GLZSM Zone Entropy, GLCM Cluster Tendency, First Order Entropy, GLDM Dependence Non-Uniformity Normalized, GLRLM Run Length Non-Uniformity* and *NGTDM Busyness)* are associated with image heterogeneity and one (*Large Area Low Gray Level Emphasis)* is associated with the proportion of large zones with low gray values within the image (9,10).

**Fig. 3:**
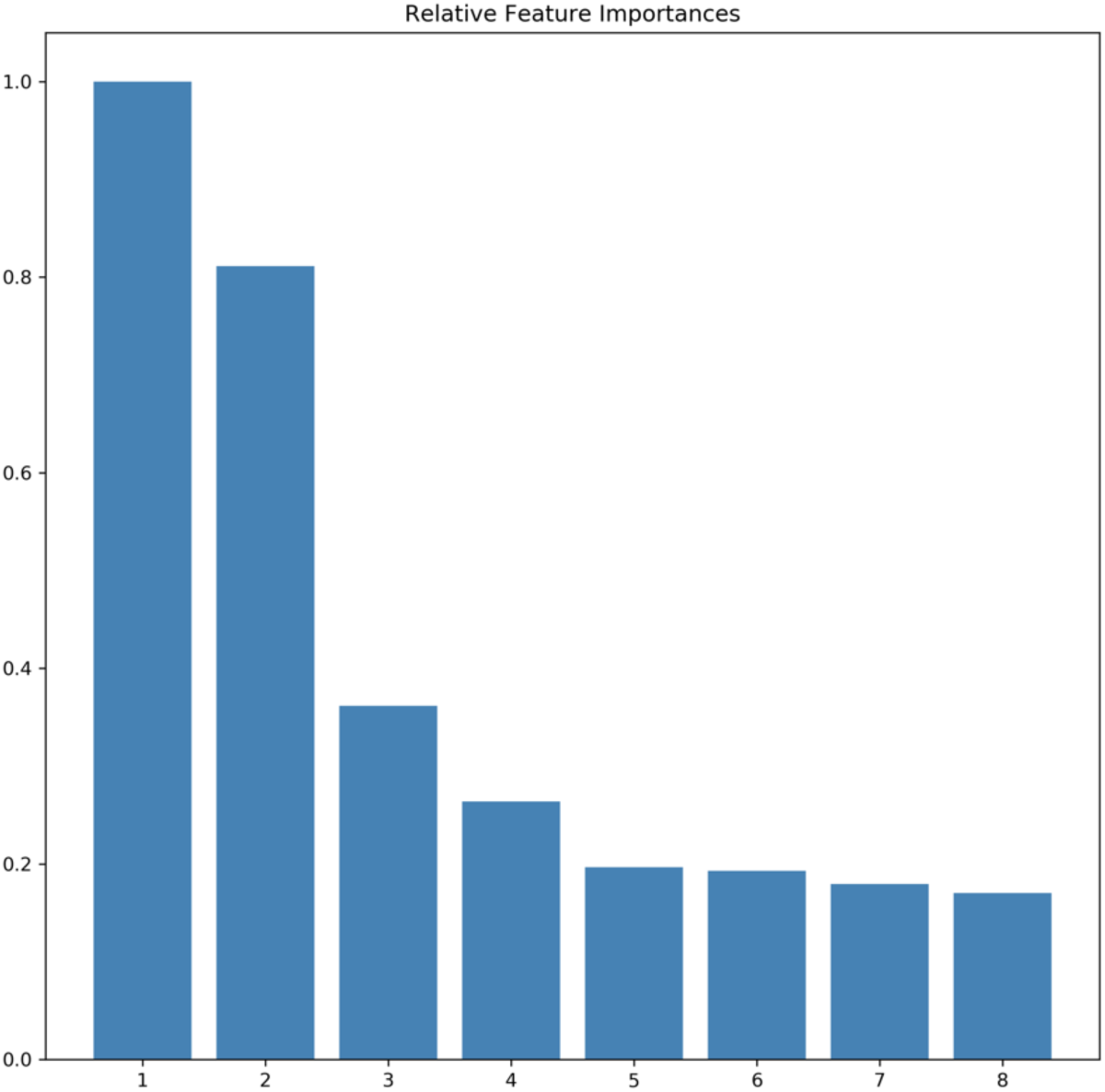
Bar plot of the 8 most important features for overall model performance as determined by the Random Forest Model. Feature importance has been normalized to the most important feature. The features, in order of descending importance are (from 1-8): *GLCM Difference Variance, GLZSM Zone Entropy, GLCM Cluster Tendency, First Order Entropy, GLDM Dependence Non-Uniformity Normalized, GLZSM Large Area Low Gray Level Emphasis, GLRLM Run Length Non-Uniformity* and *NGTDM Busyness.*

## Discussion

In this work we present a prospectively validated machine learning algorithm, which enables the prediction of overall survival and shows strong association with histopathologically defined molecular subtypes recently identified in PDAC. Several of the most important imaging features belong to a class of *heterogeneity related* features, offering explainable insights into the algorithm.

The potential of radiomics in non-invasive prediction of clinically relevant parameters, such as response to a specific therapy or expected overall survival has been shown in recent literature: For example, CT-derived radiomic signatures were shown to enable prediction of local disease control and overall survival in PDAC (11,12) or tumor grading in pancreatic neuroendocrine tumors (13). The large-scale implementation of such tools thus has the potential to become a *game changer* in medical image interpretation and individualized patient care.

*Post mortem* analyses of terminal stage PDAC specimens have shown higher tumor cellularity compared to resectable PDAC specimens, which likely represent earlier tumor development stages (14). In line with this observation we previously demonstrated that higher regional tumor cellularity identified in PDAC resection specimens was associated with a significantly worse overall survival and that the pre-operative DW-MRI-derived ADC parameter could serve as a non-invasive marker thereof (15,16). Upholding these findings, the current analysis identified the radiomic feature *Large Area Low Gray Level Emphasis*, representative of cohesive zones with low ADC values, as one of the 8 most important features for survival classification. The restriction of the pre-trained model to this single feature applied to the prediction of survival in the external validation cohort still yielded clear, albeit statistically nonsignificant separation of above and below-median predicted survival cases (see Kaplan-Meier plot and associated metrics in the supplementary material).

Several of the features ranked highly by our model *(GLCM Difference Variance, Entropy, Non-uniformity, Busyness*) represent the local heterogeneity of the image. *Entropy-*related and *Cluster Tendency* features were described in the very recent publication by Khalvati et al. as predictive of overall survival in PDAC (12). *Entropy* has furthermore been found to represent a highly reproducible and consistent imaging feature in several tumor entities and across modalities (17). The discovery of such reproducible parameters is a key part of the radiomic process and it is encouraging to see the same radiomic markers emerge not only across pancreatic cancer studies but also in other tumor entities and across different MRI systems and field strengths, supporting assumptions of overarching ontologies such as *tumor heterogeneity* and paralleling the notions of pathway-as opposed to tissue-specific therapy approaches (18).

Until proven thoroughly in large prospective trials, the inclusion of machine learning-derived predictions in a clinical decision process is ethically unjustified. However, machine-learning derived information could already be integrated into the clinical work-up of PDAC patients by back-projection of relevant radiomic features into the image space as has been demonstrated in the prostate (19) and shown in domains outside medical imaging for deep convolutional neural networks (20). Such visualizations would both aid model explainability and offer guidance for invasive tumor sampling in PDAC. The introduction of machine learning as a clinical decision support tool would also profit from the ability of machine learning algorithms to predict e.g. molecular signatures such as *KRAS* amplification status (21), that may then help stratify patients in clinical routine. Such radio-genomic approaches could complement histomorphology-derived tumor subtype prediction techniques demonstrated here, and advance the role of radiomics in precision medicine.

We selected the *Random Forest* model over the frequently-used linear models such as logistic regression for its capability of modeling both linear and non-linear relationships between features and outcomes, robustness to overfitting by design, and inbuilt insights into feature importance aiding model parsimony and explainability. *Random Forests* have also been shown to yield excellent results in previously published radiomic studies (22).

As part of any radiomic study, feature preprocessing and stability checking is required to obtain reproducible results, resulting in the majority of derived features being discarded before modeling begins (23). These discarded features are therefore rendered useless for the modeling process. To obtain more usable features, standardized acquisition and feature extraction is necessary. Recent initiatives aim to homogenize acquisition protocols between sites to enable further sequences to be included in analyses (24). We adhered to (and strongly support) the standards set by IBSI/PyRadiomics (6,10), which provide a robust post-processing platform entirely based on open-source tools, thus laying the foundation for open and reproducible radiomic science.

Our work is a proof of concept contribution to the fast-developing field of machine learning in medical imaging. Notable limitations include training cohort size, due to which the model could not reach its full potential performance (see the *training curve* in the supplemental material). The age of the imaging material in the retrospective training cohort also impacted results with several patients being excluded due to technical image quality. The quality of MRI acquisitions has since considerably improved and our results could benefit from the application of state-of-the art abdominal imaging including high resolution protocols, such as reduced field-of-view DWI (25–27). We eliminated all features that were classified unstable between the two MRI systems and recent research has provided evidence that the quantitative nature of ADC maps results in large numbers of stable features in different tumor entities and across different field strengths and MRI systems (28). Despite that, the impact of switching MRI systems between the two cohorts cannot yet be conclusively resolved. Lastly, although rigorously quality-controlled, our approach still relies on manual tumor segmentation, since recent fully-automated segmentation algorithms fail to match human observers in pancreatic tumors (29). We believe that future work will result in optimized algorithms that enable a higher level of automation -and thus standardization-of this task.

In conclusion, we show the promise of machine learning-based radiomic analyses in PDAC. We encourage the validation of the identified radiomic parameters in larger, prospectively accrued cohorts to lay the foundation for therapeutic interventions based on quantitative imaging biomarkers.

