## Supplemental Material for "A prospectively validated machine learning model for the prediction of survival and tumor subtype in pancreatic ductal adenocarcinoma"

### Supplementary Material

Georgios Kaissis[1], Sebastian Ziegelmayer[1], Fabian Lohöfer[1], Hana Algül[2], Matthias Eiber[3], Wilko Weichert[4], Roland Schmid[2], Helmut Friess[5], Ernst Rummeny[1], Donna Ankerst[6], Jens Siveke[7], Rickmer Braren[1]¶

[1]Department of Diagnostic and Interventional Radiology, Faculty of Medicine, Technical University of Munich, Munich, Germany;

[2]Department of Internal Medicine II, Faculty of Medicine, Technical University of Munich, Munich, Germany;

[3]Department of Nuclear Medicine; Faculty of Medicine, Technical University of Munich, Munich, Germany;

[4]Department of Pathology, Faculty of Medicine, Technical University of Munich, Munich, Germany;

[5]Department of Surgery, Faculty of Medicine, Technical University of Munich, Munich, Germany;

[6]Department of Mathematics, Technical University of Munich, Garching, Germany

[7]West German Cancer Center, University of Essen, Essen, Germany;

¶ Corresponding author. Author information:

Rickmer F. Braren

Attending Physician

Institute of diagnostic and interventional radiology

Faculty of Medicine

Technical University of Munich

Ismaninger Str. 22

DE-81675 Munich

Germany

#### 1 Feature extraction

Following manual tumor segmentation, the following steps were undertaken for feature extraction:

Intensity discretization was performed to a fixed bin number of 32 bins for all scans as described in (1). Due to the quantitative nature of the ADC map, no normalization was performed. Value plausibility was ascertained for all images at initial assessment. Images were spatially resampled to 3x3x3mm using the *BSpline* interpolator. No resegmentation was performed. Radiomic features were derived using PyRadiomics v. 2.1.1 (2) yielding 19 first order statistics, 16 3D shape-based, 10 2D shape based, 24 Gray Level Cooccurence Matrix, 16 Gray Level Run Length Matrix, 16 Gray Level Size Zone Matrix, 5 Neighbouring Gray Tone Difference Matrix and 14 Gray Level Dependence Matrix features as well as Laplacian of Gaussian-filtered, wavelet-decomposition-based (using the *coiflet 1* function), square, exponential, gradient, square-root, logarithm and local binary-pattern filtered versions of these features. Laplacian of Gaussian-filtering was performed with a kernel (*sigma)* of 3mm. GLCM and GLRLM were extracted using the default settings (separately for each direction then averaged). Feature descriptions can be found in the PyRadiomics documentation (3). Since radiomic feature benchmarking using the digital phantom and the included datasets (both CT) was removed in Version 7 of the IBSI manual (4) due to lack of consensus, no benchmarking was performed.

#### 2 Feature preprocessing

To ascertain comparability of radiomic features between the two MRI machines, 10 randomly selected patients from each cohort were assessed by drawing a 10mm spherical volume of interest in reference tissues (psoas muscle, gluteal fat, caudate lobe of liver). The concordance-correlation coefficient was calculated between the radiomic features derived from these measurements and features with a value below 0.8 were removed from the analysis.

Furthermore, a random sample of 20 patients each, from the training cohort and the external validation cohort, were segmented a second time by two experienced observers blinded to the initial segmentation for purposes of inter-observer stability testing. The intra-class-correlation coefficient (two-way mixed effects model/ consistency as described by McGraw and Wong (5)) was calculated and features yielding values below 0.8 were excluded.

Lastly, we excluded the following radiomic features: Features yielding nil, constant or-missing values and features with a strong correlation (*Spearman's r* >0.8*)* with tumor volume. In total, 504 reproducible and stable features were retained from 1688 features initially extracted.

**3 Feature engineering and machine learning modeling**

The following process was performed using the Python programming language version 3.7. `Monospaced` font indicates computer code that can be executed inside a Python development environment. The following packages were used: `Pandas 0.24.2, Numpy 1.16.2, Matplotlib 3.0.3, Seaborn 0.9.0, Scikit Learn 0.20.3, Lifelines 0.21.0, Statsmodels 0.9.0, Scikit Plot 0.3.7.`

Feature engineering was carried out by fitting the `sklearn.preprocessing StandardScaler` method with `feature_range=(0,1)` to the training data only and transforming both the training and the testing data with the fitted scaler.

The random forest model used was a bootstrapped `sklearn.ensemble.RandomForestClassifier.` For assessing the optimal hyperparameters, a randomized grid search was performed over the following hyperparameter settings and feature ranges:

```
grid={
'max_depth': [10, 20, 30, 40, 50, 60, 70, 80, 90, 100, None],
'min_samples_leaf': [1, 2, 4, 8, 12, 16, 20, 24, 30],
'min_samples_split': [2, 3, 4, 5, 6, 7, 8, 9, 10, 15, 20, 30],
'n_estimators': [200, 400, 600, 800, 1000, 1200, 1400, 1600, 1800,
2000, 2200, 2400, 2600],
```

```
"criterion":["gini", "entropy"]}
```

For the randomized search the `sklearn.model_selection.RandomizedSearchCV` method was used. The settings used for the randomized search were: 100 iterations and 10-fold-cross-validation resulting in 1000 model fits. The optimal parameters returned from the grid search were:

```
{'n_estimators': 2000,
 'min_samples_split': 5,
 'min_samples_leaf': 2,
 'max_depth': 30,
 'criterion': 'gini'
}
```

#### 4 Model evaluation

Model evaluation was performed using the confusion matrix and corresponding ROC curve detailed in the main text. The classification threshold was set at 0.5 (the Kolmogorov-Smirnov-statistic-derived optimal threshold was determined to be 0.49).

The cumulative recall gains curve for the testing cohort is seen below. Predicted below-median survival class (green curve, Class 1) relative gains could be maximized at 70% of the testing cohort, corresponding to 21 patients, while 80% of the population, i.e. 24 patients are required to realize the full gains for predicted above-median-survival patients as well (blue curve, Class 0).

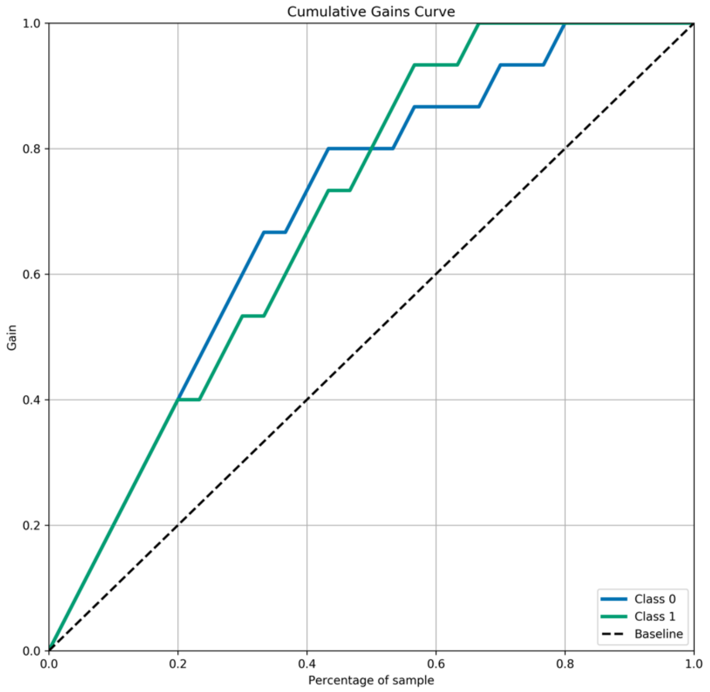


The model training curve for sensitivity is shown below. Since the statistic is derived from 10-fold cross validation, approximately 90 patients are used in every epoch to train the model. Note that both bootstrapping and cross-validation were employed to ascertain that the model encounters all training cases. No plateau is observed, indicating that model performance could have possibly been enhanced by further increasing the size of the training cohort.

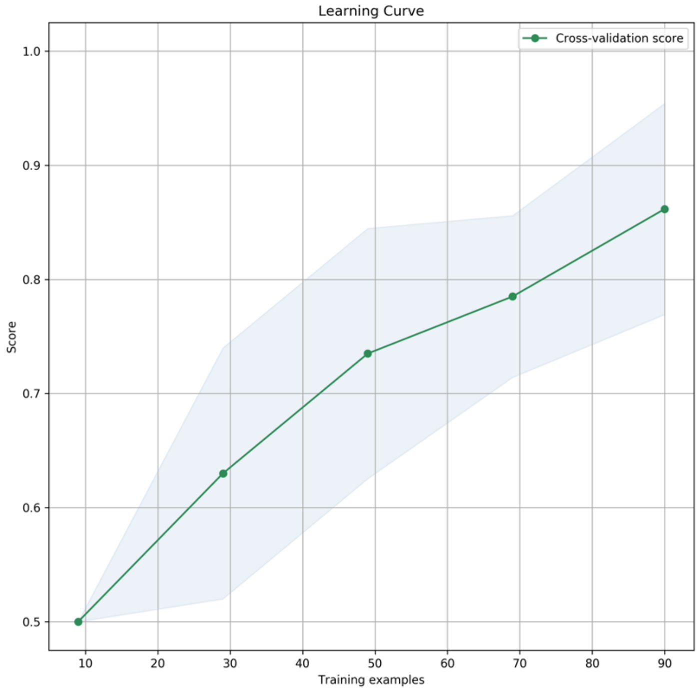


#### 4 Feature importance ranking

*Random Forests* provide inbuilt feature importance ranking by calculating which splits lead to the greatest decrease in *node impurity*, assessed by the *gini* criterion. The following graph shows the 50 most important features ranked by absolute importance:

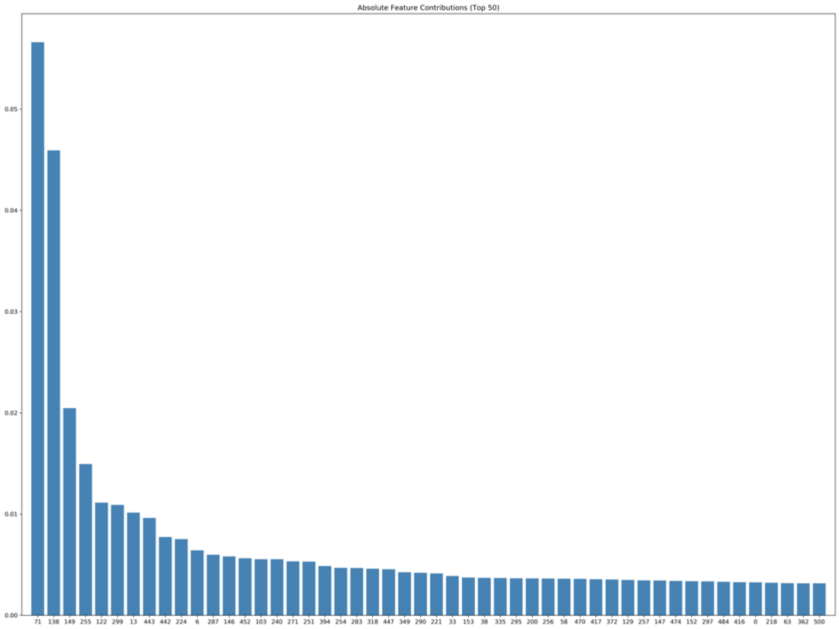

The results of recursive feature elimination using the `sklearn.feature_selection.RFECV` method employing stratified 10-fold cross validation via the `sklearn.model_selection.StratifiedKFold` method are shown below. Diminishing returns are witnessed beyond the 8 features discussed in the main manuscript.

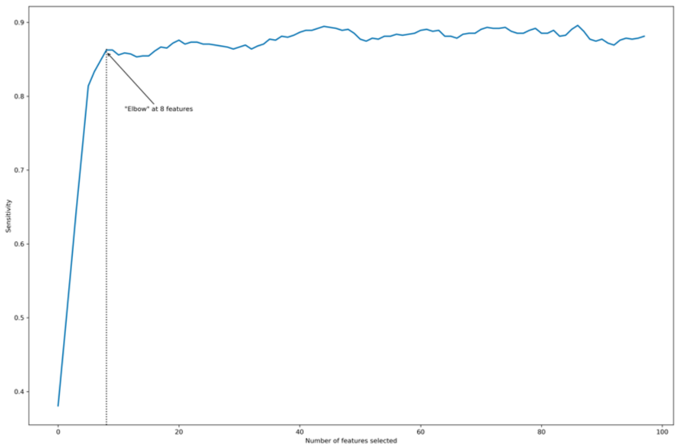


#### 5 *Large Area Low Gray Level Emphasis* assessment as a singular predictive feature

As discussed in the main manuscript, we assessed the *Large Area Low Gray Level Emphasis* feature as a singular predictor of above vs. below median survival in the external validation cohort. The model suffered a sharp loss in sensitivity (53%) but still achieved a specificity of 79% in the external validation set. The Kaplan Meier plot below shows the hypothetical Kaplan-Meier curves of the patients as predicted by the model. No statistical significance is observed and there is violation of the proportional hazards assumption at early timepoints (*log-rank-test* P=0.13).

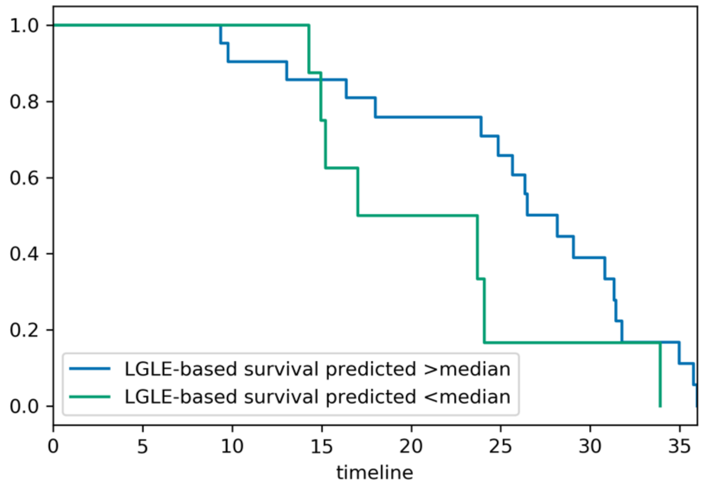


#### 6 Results of clinical parameter cross-tabulation and multivariate proportional hazards survival modeling

To exclude any confounding effect from clinical parameters, we investigated their impact on overall survival and their distribution between the two groups as detailed in the main manuscript. In brief, all confounding clinical parameters were equivalently distributed between groups. In the training cohort, choice of chemotherapy regimen and metastatic state were significantly associated with survival time. In the external validation cohort, only metastatic state was significantly associated with survival time.

Multivariate Cox-regression results are shown below.

pT was treated as an ordinal variable. LNR was stratified as <0.2 including pN0 vs. ≥0.2 according to (6). Groupings for p/cN, p/cM, G and R and ECOG result from available groups.

| Parameter | P | HR | 95.0% CI for HR | |
|---|---|---|---|---|
| | | | Lower | Upper |
| p/cT (2-3-4) | .198 | 1.203 | .908 | 1.595 |
| p/cN (0 vs. 1) | .350 | 1.524 | .630 | 3.682 |
| G (2 vs. 3) | .208 | 1.351 | .846 | 2.159 |
| R (0 vs.1) | .711 | 1.095 | .676 | 1.776 |
| p/cM (0 vs. 1) | .030 | 1.871 | 1.303 | 2.453 |
| LNR | .070 | 1.522 | 1.025 | 1.923 |

| | | | | |
|---|---|---|---|---|
| Age | .284 | .992 | .977 | 1.007 |
| Sex | .911 | .975 | .630 | 1.509 |
| Tumor Volume | .563 | .992 | .966 | 1.019 |
| ECOG (0 vs. 1) | .802 | .949 | .821 | 1.202 |
| CTX (Gem. vs. FOLFIRINOX) | .007 | 1.996 | 1.211 | 3.289 |

Multivariate Cox Regression for survival times in the training cohort.

| Parameter | P | HR | 95.0% CI for HR | |
|---|---|---|---|---|
| | | | Lower | Upper |
| p/cT (2-3-4) | .929 | .965 | .438 | 2.125 |
| p/cN (0 vs. 1) | .349 | .896 | .523 | 3.789 |
| G (2 vs. 3) | .380 | 1.815 | .479 | 6.877 |
| R (0 vs. 1) | .846 | .882 | .247 | 3.148 |
| p/cM (0 vs. 1) | .027 | 2.003 | 1.672 | 2.677 |
| LNR | .190 | 1.09 | .670 | 1.408 |
| Age | .074 | .937 | .872 | 1.006 |
| Sex | .958 | 1.036 | .284 | 3.774 |
| Tumor Volume | .200 | .965 | .914 | 1.019 |
| ECOG (0 vs. 1) | .775 | 1.010 | .702 | 1.364 |
| CTX (Gem. vs. FOLFIRINOX) | .655 | .717 | .166 | 3.088 |

Multivariate Cox Regression for survival times in the external validation cohort.

The distribution of clinical data between the two cohorts is shown below. None of the parameters was significantly overrepresented between the training and external validation cohorts.

| Parameter | Training Cohort | STDEV | Testing Cohort | STDEV | P |
|---|---|---|---|---|---|
| **Age** | 60.7 | 15.3 | 61.4 | 8.78 | |
| **Chemotherapy regimen** | Gem: 70.6% (72)<br>FOLFIRINOX: 29.4% (30) | | Gem: 70% (21)<br>FOLFIRINOX: 30% (9) | | 1.0 |
| **Experienced Event** | Yes: 90.2% (92)<br>No: 9.8% (10) | | Yes: 86.7% (26)<br>No: 13.3% (4) | | .52 |
| **G** | 2: 55.9% (57)<br>3: 44.1% (45) | | 2: 56.7% (17)<br>3: 43.3% (13) | | 1.0 |
| **Overall Survival (Months)** | 20.5 | 7.6 | 23.4 | 8.0 | .07 |
| **p/cM** | 0: 77 (75%)<br>1: (25%) | | 0: 22 (73%)<br>1:8 (27%) | | .81 |
| **p/cN** | 0: 38.2% (39)<br>1: 61.8% (63) | | 0: 46.7% (14)<br>1: 53.5% (16) | | .52 |
| **p/cT** | 2: 26.5% (27)<br>3: 36.3% (37)<br>4: 37.3% (38) | | 2: 26.7% (8)<br>3: 36.7% (11)<br>4: 36.7% (11) | | (1.0) |
| **R** | 0: 56.9% (58)<br>1: 43.1% (44) | | 0: 53.3% (16)<br>1: 46.7% (14) | | .83 |
| **Sex** | Female: 45.1% (46)<br>Male: 54.9% (56) | | Female: 46.7% (14)<br>Male: 53.3% (16) | | 1.0 |
| **Tumor Volume (ml)** | 18.08 | 8.47 | 20.02 | 8.91 | .27 |
| **ECOG** | 0: 82 (80%)<br>1: 20 (20%) | | 0: 22 (73%)<br>1: 8 (27%) | | .45 |
| **Lymph Node Ratio** | .11 | .08 | .09 | .10 | .24 |

Distributions of clinical parameters between the groups. Averages and standard deviations are given (where applicable). P-values refer to *Chi-squared/Fisher's exact tests* for crosstabulation.

Pre-operative levels of the CA 19-9 tumor marker were available for 33 patients in the training cohort with an average value of 365.4 [5 to 2360] U/ml and for 8 patients in the external validation cohort with an average value of 369.3 [36 to 2370] U/ml (unpaired t-Test P= 0.98).

#### 7 STROBE Statement Checklist

Data collection and processing is reported according to STROBE guidelines. The checklist with the pertinent locations of the reported data is provided below

| | Item No | Reccomendation | Remark/ Location |
|---|---|---|---|
| **Title and abstract** | 1 | (*a*) Indicate the study's design with a commonly used term in the title or the abstract | Reported in title |
| | | (*b*) Provide in the abstract an informative and balanced summary of what was done and what was found | Reported in abstract |
| **Introduction** | | | |
| Background/rationale | 2 | Explain the scientific background and rationale for the investigation being reported | Introduction §§ 1-3 |
| Objectives | 3 | State specific objectives, including any prespecified hypotheses | Introduction § 3 |
| **Methods** | | | |
| Study design | 4 | Present key elements of study design early in the paper | Mehods/ Study Design |
| Setting | 5 | Describe the setting, locations, and relevant dates, including periods of recruitment, exposure, follow-up, and data collection | Ibid. |
| Participants | 6 | (*a*) Give the eligibility criteria, and the sources and methods of selection of participants. Describe methods of follow-up | Ibid. |
| | | (*b*) For matched studies, give matching criteria and number of exposed and unexposed | N.A. |
| Variables | 7 | Clearly define all outcomes, exposures, predictors, potential confounders, and effect modifiers. Give diagnostic criteria, if applicable | Ibid. and Methods/ Clinical variables. Compare also Results §§ 4-5 |
| Data sources/ measurement | 8* | For each variable of interest, give sources of data and details of methods of assessment (measurement). Describe comparability of assessment methods if there is more than one group | Methods/ Study Design |
| Bias | 9 | Describe any efforts to address potential sources of bias | Supplementary Material §2, Results §§4-5 |
| Study size | 10 | Explain how the study size was arrived at | Methods/ Study Design |
| Quantitative variables | 11 | Explain how quantitative variables were handled in the analyses. If applicable, describe which groupings were chosen and why | Results §4 |
| Statistical methods | 12 | (*a*) Describe all statistical methods, including those used to control for confounding | Methods/ Inferential Statistical Modeling |
| | | (*b*) Describe any methods used to examine subgroups and interactions | Ibid. |
| | | (*c*) Explain how missing data were addressed | Supplementary Material §2 Results §5. |

| | | | |
|---|---|---|---|
| | | (*d*) If applicable, explain how loss to follow-up was addressed | Methods/ Study Design |
| | | (*e*) Describe any sensitivity analyses | Supplementary Material/ §4 |
| **Results** | | | |
| Participants | 13* | (a) Report numbers of individuals at each stage of study—eg numbers potentially eligible, examined for eligibility, confirmed eligible, included in the study, completing follow-up, and analysed | Methods/ Study Design Compare Flow-Charts below |
| | | (b) Give reasons for non-participation at each stage | Ibid. |
| | | (c) Consider use of a flow diagram | See below |
| Descriptive data | 14* | (a) Give characteristics of study participants (eg demographic, clinical, social) and information on exposures and potential confounders | Methods/ Study design Results §§4/5 |
| | | (b) Indicate number of participants with missing data for each variable of interest | Results §5 |
| | | (c) Summarise follow-up time (eg, average and total amount) | Methods/ Study design |
| Outcome data | 15* | Report numbers of outcome events or summary measures over time | Results §2 and 4/5 |
| Main results | 16 | (*a*) Give unadjusted estimates and, if applicable, confounder-adjusted estimates and their precision (eg, 95% confidence interval). Make clear which confounders were adjusted for and why they were included | Results §4/5 |
| | | (*b*) Report category boundaries when continuous variables were categorized | N.A. |
| | | (*c*) If relevant, consider translating estimates of relative risk into absolute risk for a meaningful time period | N.A. |
| Other analyses | 17 | Report other analyses done—eg analyses of subgroups and interactions, and sensitivity analyses | N.A. |
| **Discussion** | | | |
| Key results | 18 | Summarise key results with reference to study objectives | Discussion §1 |
| Limitations | 19 | Discuss limitations of the study, taking into account sources of potential bias or imprecision. Discuss both direction and magnitude of any potential bias | Discussion §7 |
| Interpretation | 20 | Give a cautious overall interpretation of results considering objectives, limitations, multiplicity of analyses, results from similar studies, and other relevant evidence | Discussion §§ 2-6 |
| Generalisability | 21 | Discuss the generalisability (external validity) of the study results | Introduction §4 Discussion §§ 2, 4, 7 Supplementary Material §4 |
| **Other information** | | | |
| Funding | 22 | Give the source of funding and the role of the funders for the present study and, if applicable, for the original study on which the present article is based | Preamble |

Flow charts of patient recruitment for the training and testing cohorts are shown below:

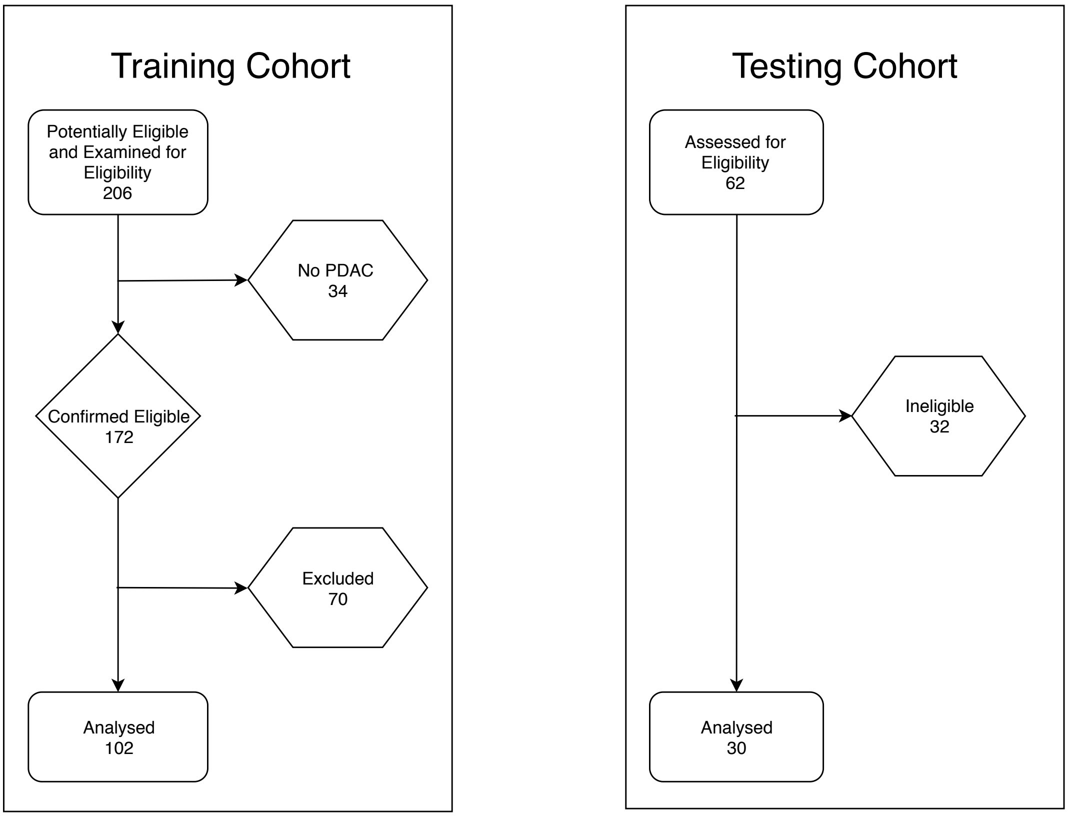


Reasons for exclusion:

-Training cohort: No PDAC in final histopathological report (34 patients), no mass detectable in imaging (3 patients), insufficient technical quality of imaging (24 patients), study aborted by patient (4 patients), no DWI performed due to incompatible cardiac pacemaker (1 patient), death within 2 months of imaging (21 patients), received prior treatment or refused treatment (17 patients)

-Testing cohort: Received prior treatment or refused treatment or study inclusion (17 patients), no PDAC in final histopathological diagnosis (12 patients), insufficient technical quality of imaging (2 patients), died within 2 months of imaging (1 patient).

*Compare also*: https://pyradiomics.readthedocs.io/

(3) PyRadiomics feature definitions:

https://pyradiomics.readthedocs.io/en/latest/features.html

*Compare also*: (4) below

(4) Zwanenburg, A., Leger, S., Vallières, M., and Löck, S. (2016). Image biomarker standardisation initiative: https://arxiv.org/abs/1612.07003v7

Multiple versions of the manual are available, version 7 is linked and was current at the time of the analysis

(5) McGraw, K. O., & Wong, S. P. (1996). Forming inferences about some intraclass correlation coefficients. *Psychological Methods, 1*(1), 30-46.

(6) Bilici A. (2014). Prognostic factors related with survival in patients with pancreatic adenocarcinoma. *World J Gastroenterology*, 20(31): 10802–10812.
